# A Mathematical Framework for Understanding Recognition Systems

**DOI:** 10.1101/2023.06.08.544240

**Authors:** Yusuf Brima

## Abstract

Recognition plays a crucial role in the formation of social bonds, mate selection, kin selection and survival across species. However, a unifying framework for understanding recognition systems in biology is lacking. We, therefore, propose a theoretical framework for biological recognition utilizing basic principles from category theory, information theory, dynamical systems modeling and optimization theory. We define the two types of recognition, individual recognition (IR) and class-level recognition (CR), as *functors* between categories of *stimuli* and *responses*. IR produces *unique* responses for each individual, while CR produces *shared* responses for multiple individuals of the same class. We identify five conditions – *universality, low entropy, unfalsifiability, uniform convergence* and *cognitive limit* – that must hold for robust IR systems, which we term “signature systems.” Further, we model signature systems as *attractor states* with perspectives from both statistical information processing and dynamical systems. Our framework provides a basis for advancing understanding of the mechanisms underlying biological recognition and its implications for communication and behavior. Overall, the mathematical conceptualization of IR provides a basis for advancing our understanding of communication and the evolution of language.

## 1 Introduction

Recognition plays a crucial role in an animal’s ability to identify and distinguish between different objects, (this could include living things like self or other individuals) based on their characteristics. This ability is particularly significant for the survival and evolution of animals under evolutionary pressures. Animals rely on recognition to navigate their environment, identify conspecifics, and avoid predators, enabling them to engage in complex social behaviors [1, 2]. To achieve this feat, social animals have evolved and fine-tuned their sensory systems, which have been extensively studied in the context of animal communication and behavior. However, despite significant research efforts, a well-established mathematical framework for recognition in this domain is lacking, and there is no unifying framework to describe the various recognition systems and their properties [3–7].

The study of recognition systems (definition is stated in Section 2) has traditionally been focused on understanding its ecological and evolutionary significance [8–10]. However, recent research has highlighted the need to develop a formalism for recognition in biology, particularly to understand the mechanisms and processes underlying these systems [1, 8–10]. We refer to formalism as the use of mathematical (and computational) models to describe and analyze complex systems, a case in point, recognition systems in biology. The development of a formalism for recognition in biology could potentially allow for a more systematic and rigorous analysis of these systems, enabling us to identify commonalities and differences within and across species and to test hypotheses about the mechanisms underlying recognition.

In this context, category theory, dynamical systems modeling, optimization theory and information theory have emerged as promising approaches for developing models of systems and processes either artificial or biological (or a combination of both). Category theory provides a general framework for describing and analyzing complex systems, including those found in biology. It has been successfully applied in other areas of biological sciences, such as neuroscience and genetics, to provide a formal language for describing complex systems and their interactions [11–15]. Moreover, biology as an evolutionary process fine-grainly fits in the framework of dynamical systems model. This could include, among other things, dynamics of a signature (unique cues that identify objects in a general sense) evolution of individuals in a signature system, predator-prey interaction, mate selection dynamics, etc. Optimization theory on the other hand, is suitable for modeling recognition from the perspective of a receiver, for example, in an abstracted sense, optimally finding the identity of each learned individual identity in every instantiation of meetings or environmental configurations. Underlying all this process is the information-theoretic nature of the signaling systems, that is, the cue production by the signaler, propagation through a medium, perception by a receiver, etc. Every step in the way involves deep (by deep we mean hierarchical) information processing that is constrained by many factors (such as cue production, dominance hierarchy of the signaler in their social ladder, health status, kind of interact whether predatory or not), medium of transmission, perceptual processing quality and many more factors of variability. Most of this information-theoretic formulation holds true from the perspective of the signaler as well, such as the production quality of cues, perceiving and interpreting receiver response which all could cascade further processes such as Theory of Mind of the receiver, for example.

As animal behavior and communication involves signaling either intentionally or unintentionally, which from a signal processing standpoint involves Information Theory. We, therefore, utilize elements of Information Theory to explain this phenomenon. So, we hope the application of these mathematical tools would have the potential to provide a unifying framework for describing the various types of recognition, such as Individual Recognition (IR) and class-level recognition, and their underlying mechanisms.

The remaining sections of this paper is organized as follows: we present a brief overview of recognition systems in Section 2. In Section 3, we propose of a theoretical framework for understanding recognition systems based on mathematical principles from category theory. This leads to review IR as a mathematical optimization problem in Section 4, and signature systems as the bases for IR are discussed in Section 5 where we present conditions that must hold for signature systems to ensure reliable IR. In Section 6, we show that signature systems to attractor states, and in Section 7, we discuss of potential implications of the our proposed framework. Finally, we end with concluding remarks in Section 8.

## 2 Recognition Systems

Recognition systems, in the context of biology, refer to integrated mechanisms that gives an individual the ability to distinguish between different individuals or objects based on certain cues or features. These cues can be visual, auditory, olfactory, or tactile in nature. We have shown a high-level view of this integrated mechanisms in Fig. 1.

**Fig 1.**
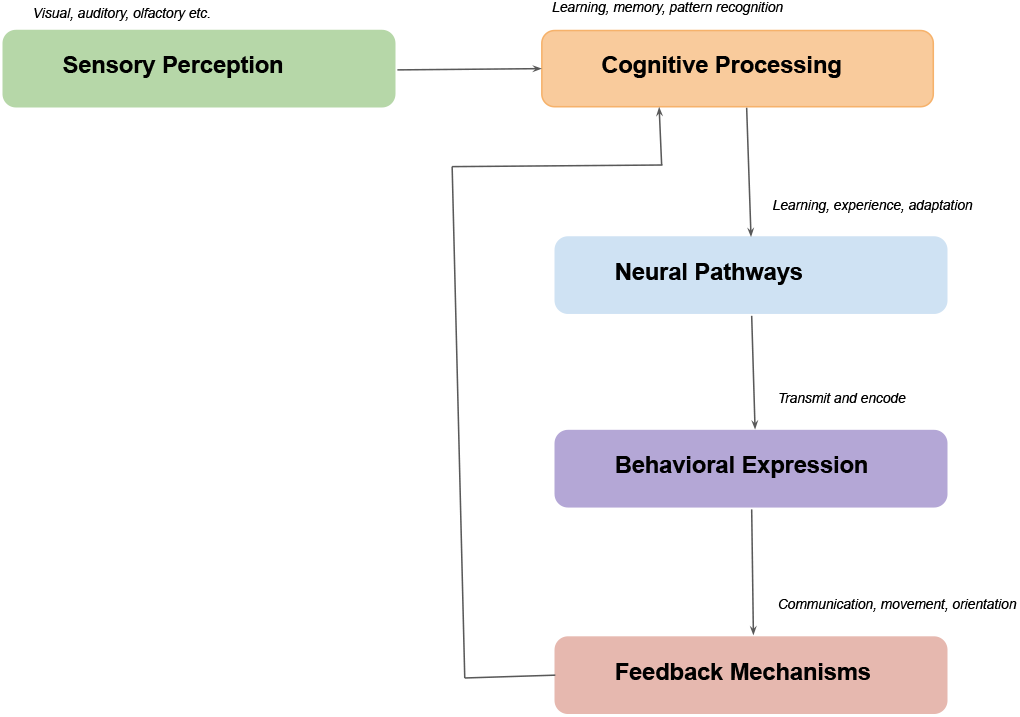
The process of recognition from the perspective of a receiver involving sensory perception through cognitive mechanisms to behavioral response and feedback optimization. Sensory perception detects cues which are processed cognitively through learning, memory, and pattern recognition. This information is transmitted through neural pathways to produce behavior. Feedback from behavior and the environment then optimizes the system.

The integrated components of a recognition system include:

- **Sensory perception**: The ability to detect cues, signals and markers that can be used for recognition. This includes visual, auditory, olfactory, tactile or chemosensory. Different species rely on different sensory modalities depending on their evolutionary adaptations.
- **Cognitive processing**: The ability to learn, remember, and categorize sensory information in meaningful ways. This includes pattern recognition, the formation of perceptual categories, spatial memory, and the association of cues with rewards, punishments or other experiences. Which all have a crucial role in the fitness of the recognizer.
- **Neural pathways**: The neural circuits and pathways that connect sensory perception to cognitive processing centers in the brain. These pathways transmit and encode information in a way that can be integrated and acted upon. Different forms of recognition, e.g. individual vs. species, involve distinct neural pathways [16, 17].
- **Behavioral expression**: The expression of recognition through communication, movement, orientation, aggression, courtship, alarm calls or other behaviors. These behaviors demonstrate that recognition has occurred and allow the recognition to impact the receiver’s interactions with the environment.
- **Feedback mechanisms**: The mechanisms by which the performance of recognition is optimized over time through learning, experience, and evolution. This includes associative learning, imprinting, habituation, and adaptation. Feedback serves to strengthen neural pathways involved in effective recognition and prune those involved in less effective recognition.

However, the focus of this present paper is not to delve into the integrated processes on an individual basis, but to create a unifying framework of this subject using mathematical constructs. In that regard, the definition of a recognition system can be abstracted as it is more general but the object of this paper is recognition in biology. So, there are two main types of recognition systems that will be explored here: individual recognition (IR) and class-level recognition (CR). IR involves the ability of an individual to differentiate between specific individuals based on *unique* and *distinctive* cues or features, such as facial patterns (we humans are really excel at this [18]) or vocalizations (our extant relatives, chimpanzees can infer individual identity from long-distance pant-hoot calls for example [19, 20]). Class-level recognition, on the other hand, involves the ability to classify individuals into *categories* based on *shared characteristics*, such as species or sex [1, 10]. Recognition systems are crucial for a variety of biological processes, including social interactions, mating behavior, and kin recognition. Examples of animals with IR systems include honeybee (*Apis*), zebrafish (*Danio rerio*), and African elephant (*Loxodonta*), while class-level recognition systems are found in animals such as primates (*Primate*) and birds (*Aves*) [1].

## 3 A Category Theory view of Recognition Systems

In this paper, we propose a formal framework that differentiates between IR and CR in recognition systems using insights from category theory, which is a branch of mathematics that provides a framework for understanding the relationships between different mathematical structures. We model recognition systems as *functors* between categories, where categories represent the set of *stimuli* and the set of *responses*. The category of stimuli refers to the stream of input signal that the recognition system receives and processes in order to produce a response. These stimuli can be of any modality or a combination thereof, such as visual, auditory, or olfactory. For example, in a face recognition system, the stimuli could be a set of images of human faces, while in a vocal recognition system, the stimuli could be a set of vocalizations or audio recordings of speech. The category of stimuli is an important concept, as it allows us to represent the inputs of a recognition system as *objects in a category*, which can be related to each other through *morphisms*, such as functors. And the category of response refers to the set of possible outputs that a recognition system can generate in response to a given stimulus. For instance, in a face recognition system, the category of responses might consist of unique identification labels that correspond to specific individuals or gender whose faces have been previously registered in the system. Similarly, in a speaker recognition system, the category of responses might consist speaker identities or gender. The response category is also a key component of recognition systems as it defines the criteria by which the system determines whether a given stimulus matches a particular response. To illustrate these concept, we have provided in Fig. 2 the two types of recognition, IR and CR, which are modeled as functors from the category of stimuli to the category of responses.

**Fig 2.**
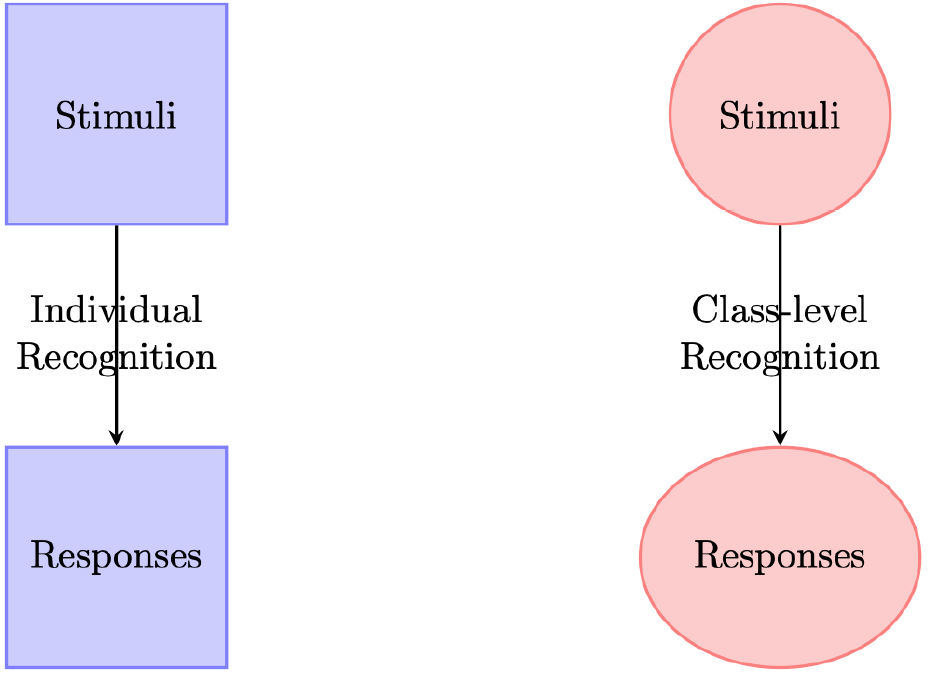
A visual illustration of the difference between IR (left) and Class-level recognition (right).

Formally, we represent IR as a functor *I* from the category of stimuli *S* to the category of responses *R*, where *I*(*s*) is the response to stimulus *s*. In this case, the responses are *unique* to each individual and cannot be generalized to other individuals. We represent this as:

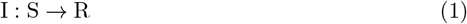

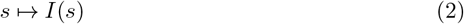

CR, on the other hand, is represented as a functor *C* from the category of stimuli *S* to the category of responses *R*, where *C*(*s*) is the response to stimulus *s*. In this case, the responses are *shared* by multiple individuals belonging to the same class. This is represented as:

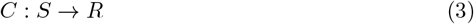

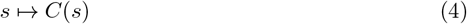

Both IR and CR satisfy the properties of category theory, specifically the properties of identity and composition. The identity property states that for every object in a category, there exists an identity morphism that maps the object to itself. In the context of recognition systems, this means that *there exists an identity recognition system id*_*S*_, *for every stimuli category S, such that id*_*S*_ *maps each stimulus s* ∈ *S to itself* as shown in Eq (5). For example, the identity recognition system for a face recognition task would map each face to itself.

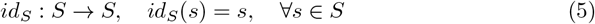

Also, the composition property states that for any two morphisms *f* and *g* in a category, there exists a morphism *h* that maps the source of *f* to the target of *g*. In this context, this means that given two recognition systems, *F* and *G*, there exists a recognition system *H* that maps the stimuli to the responses of *G*, and then maps those responses to the responses of *F*.This is represented as

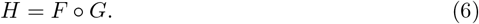

An example of the composition property could be multi-modal recognition (face and voice). Let us say we have two recognition systems, *F* for faces and *G* for voices. *F* takes a face image as input and outputs a recognition score or ID that corresponds to a particular individual’s face. Similarly, *G* takes an audio sample of a person’s voice and outputs a recognition score or ID that corresponds to that individual’s voice. Now, let’s say we want to create a recognition system that combines both face and voice recognition. We can do this by using the composition property of category theory. We can create a new recognition system *H* that takes an input of both a face image and an audio sample, uses *G* to recognize the voice and *F* to recognize the face, and then combines the two recognition scores to output a final recognition score or ID. Mathematically, *H* can be represented as Eq (6), where *G* maps the stimuli (the audio sample) to the responses (the voice recognition score) and *F* maps the stimuli (the face image) to the responses (the face recognition score).

The main difference between the two recognition systems is the *nature of the response*. IR produces *unique responses* for each individual, while CR produces *shared responses* for multiple individuals belonging to the same class. One example of a system that employs IR is the vocal recognition system in bottlenose dolphins (*Tursiops*). Each dolphin has a unique vocal signature that is used for IR [21]. In contrast, the coloration patterns in zebras (*Equus quagga*) are used for class-level recognition, where the unique stripe patterns are shared by multiple individuals belonging to the same class [22].

## 4 Individual Recognition

Individual Recognition, as the ability of an individual to distinguish between different members of its own species based on specific traits, is important for a variety of social behaviors such as mating, aggression, and cooperation. The recognition of individuals has been extensively studied in a wide range of taxa, including insects, birds, mammals, and fish [23–30]. In insects, IR can be based on olfactory cues, visual cues, or a combination of both. For example, honeybees (*Apis*) use pheromones (chemical substances which are secreted to the outside by an individual and received by a another individual of the same species) to recognize nest mates from non-nest mates [31], while paper wasps use facial patterns for IR [9]. Birds on the other hand use visual and auditory cues for IR, such as the unique songs of male birds [32]. Mammals are also capable of IR, with some species relying on olfactory cues, while others use visual or auditory cues. For example, elephants can recognize the scent of individual family members, even after prolong periods of separation [33]. In primates, facial recognition plays an important role in social interactions, with some species exhibiting the ability to recognize individual faces even in photographs [34]. Fish also use a variety of sensory cues for IR, such as olfactory, visual, and auditory cues. The African cichlid fish (*Cichlidae*) uses the coloration of males to identify individual conspecifics [35], while zebrafish (*Danio rerio*) use both olfactory and visual cues for IR [35]. Overall, IR is an important aspect of social behavior in many species and can have important implications for mating, social hierarchy, and cooperation. We have presented a high-level dyadic framework of IR in Fig 3.

**Fig 3.**
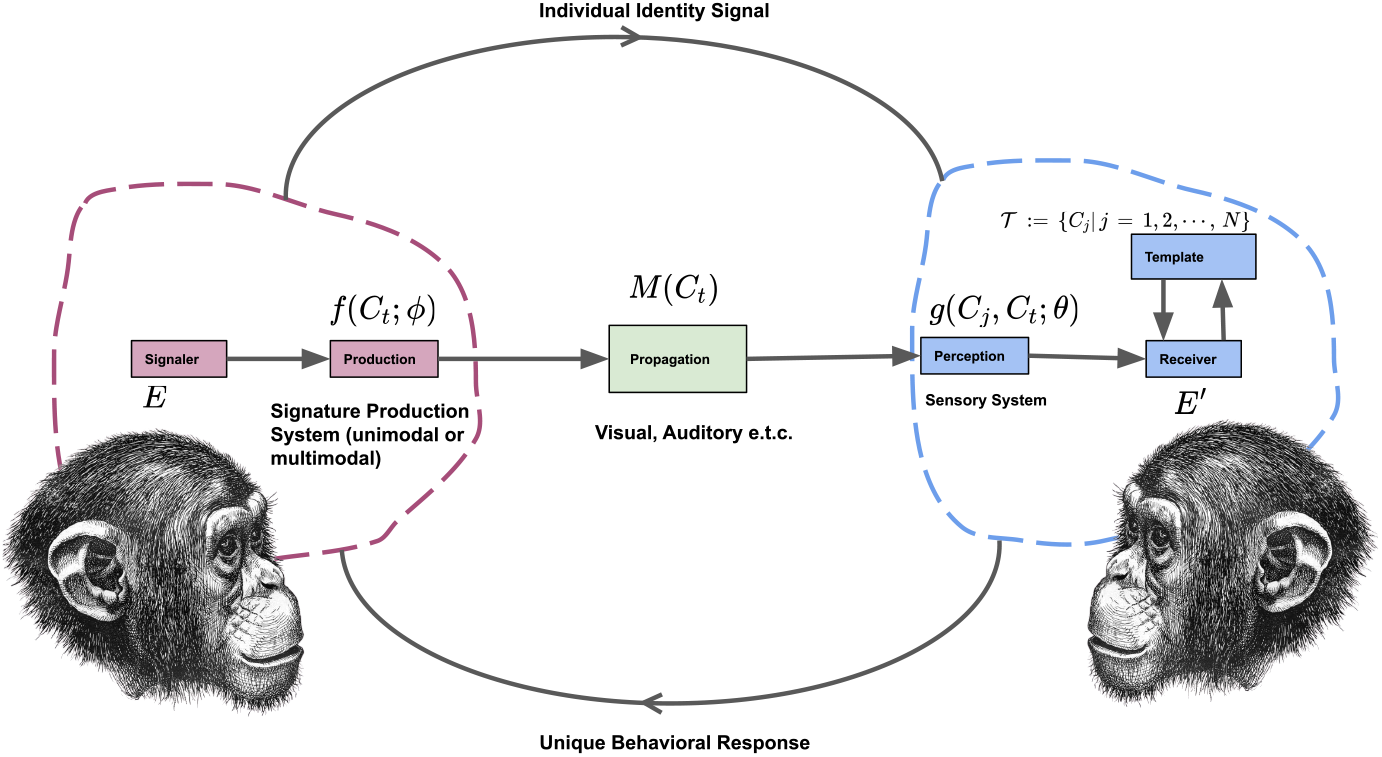
A signaler *E* at time *t* produces *f* (*C*_*t*_; *ϕ*) individually distinct cues *C*_*t*_ such as visual, acoustic or olfactory that is propagated in a medium *M* (*C*_*t*_). A cue can be uni-modal or multi-modal. A receiver *E*^*′*^ perceives *g*(*C*_*j*_, *C*_*t*_; *θ*) cues through its sensory percepts that is matched against a templating system. A cue is compared with known templates in *𝒯* to determine a unique behavioral response directed to the signaler as shown in equation (8).

The goal of IR is:

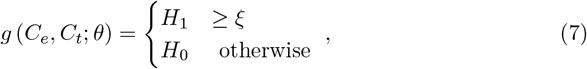

where *g* (*C*_*e*_, *C*_*t*_; *θ*) is a template matching function which takes *C*_*e*_ and *C*_*t*_, the enrolled and perceived (at time *t*) cues respectively. This function is parameterized by *θ* that encapsulates the notion of template learning for each unique individual by the receiver. In this formulation, *H*_1_ is the hypothesis that a signaler’s cue truly claims who they are while *H*_0_ is the contrary matching hypothesis given a specified acceptable cue verifiability threshold *ξ*. We have shown this matching process in Eq (8).

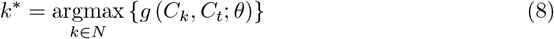

Another framing of this task is to define it as a function:

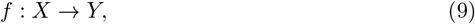

where *X* is the set of all possible individuals and *Y* is the set of all possible individual identities. The function *f* maps an individuals *x* ∈ *X* to the corresponding identity *y* ∈ *Y* such that *f* (*x*) = *y*. For IR to be successful, the function *f* must be able to distinguish between individuals with a high degree of accuracy which in equation (7) is a higher threshold *ξ*. We can express this as a measure of similarity between individuals identity cues as shown in the biological instantiation in Eq (8). Put another way, let *s*(*x, y*) be a similarity function that measures the degree of similarity between an individual *x* and an identity *y*. A successful IR system must ensure that the similarity between individuals and the corresponding identities is high, and the similarity between individuals and other identities is low.

A commonly used computational framework for IR is the pattern recognition framework. In this framing, we can view IR as a classification problem, where the goal is to classify individuals into their corresponding identities. We define IR in the pattern recognition framework as follows:

Let *X* be the *feature space*, that is, a set of all possible characteristics (cues or features) that contribute to the formation of the unique identity of an individual and *Y* be the *label space*, the set of all possible identities there exist. Given a set of sample data *{* (*x*_1_, *y*_1_), (*x*_2_, *y*_2_) *}*, …, (*x*_*n*_, *y*_*n*_), where *x*_*i*_ ∈ *X* and *y*_*i*_ ∈ *Y*, the goal of IR is to learn a mapping function *f* in Eq (9) that minimizes the expected loss over the joint distribution *P*_*X,Y*_ (*x, y*), such that:

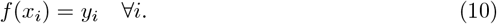

In this framework, the mapping function *f* is typically learned using machine learning algorithms, such as decision trees, hidden Markov model (HMM), linear discriminant analysis (LDA), neural networks, support vector machines (SVM) [19, 36–40]. As presented here, the choice of algorithm depends on the specific IR task and the characteristics of the data.

## 5 Signature Systems

In this section, we discuss the elements give rise to robust recognition: signature systems. Signature systems are a type of IR systems in that rely on the unique features of an individual’s characteristics or behavior. Conceptually, signature systems are a subset of recognition systems with focus on *uniqueness and distinctiveness of cues or features*. These systems are widely observed in a variety of species such as primates, birds, and insects [1, 2, 41]. An example of a signature system is facial recognition used by primates. Studies have shown that primates can distinguish between individuals based on facial features such as the shape and size of the nose, mouth, and eyes [42]. Similarly, birds use distinctive vocalizations as a signature system to identify individuals within their social groups [41]. Insects also use signature systems, such as the unique chemical signals produced by ants and bees to distinguish between individuals within their colonies [1, 2]. These signature systems have been found to play a critical role in social organization and communication in these species.

The study of signature systems in biology has led to a deeper understanding of the mechanisms of IR and the evolution of social behavior in a variety of species. By examining the unique features and signals used by different individuals, insights can be gained into the complex processes of communication and social organization in the natural world. We shown shown illustrations of four species with IR abilities in Fig. 4.

**Fig 4.**
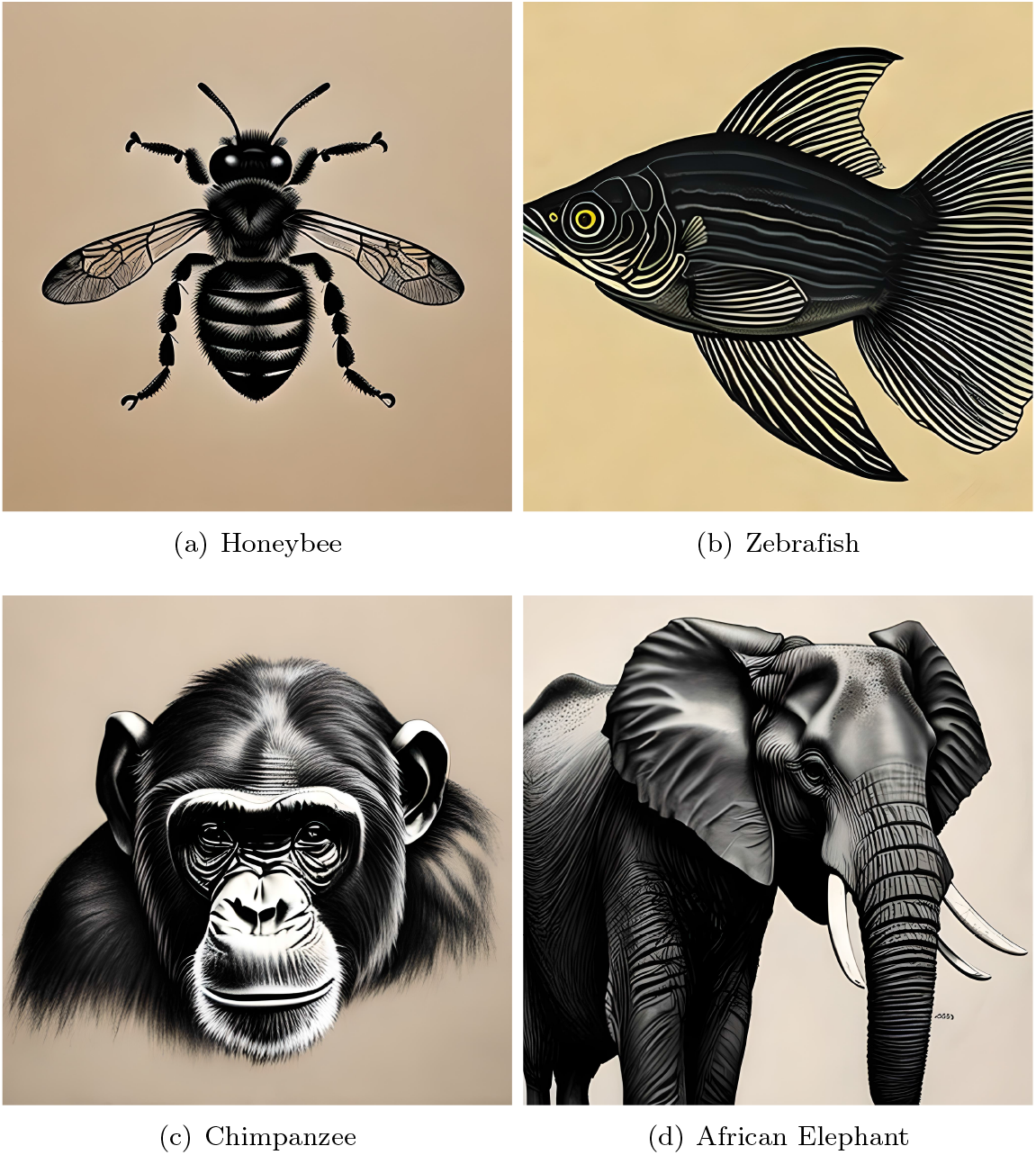
IR mechanisms used by honeybees, zebrafish, chimpanzees, and African elephants include olfactory cues, visual cues, facial recognition, and vocalizations.

From Fig. 4, honeybees use olfactory cues for IR. They have a highly developed sense of smell that enables them to detect the specific pheromones of each member of their colony [43, 44]. Each bee has a unique blend of pheromones that is influenced by factors such as genetics, age, and their environment [45, 46]. By detecting and remembering the specific pheromone blend of each individual, honeybees are able to distinguish between nestmates and non-nestmates and regulate the social behavior within the colony [47]. Zebrafish use visual cues for IR [48, 49]. They have the ability to recognize the unique pattern of stripes on the body of each member of their shoal. This pattern is determined by genetic factors and remains *stable* throughout the life of the fish [49]. By recognizing the specific stripe pattern of each individual, zebrafish are able to form stable social groups and avoid aggressive interactions with unfamiliar fish [50]. Chimpanzees use a combination of visual and auditory cues for IR [51–53]. They have the ability to recognize the unique facial features and vocalizations of each member of their social group. Facial recognition is based on features such as the shape, color, and texture of facial hair, while vocal recognition is based on the unique pitch, rhythm, and tone of an individual’s vocalizations [54–56]. By recognizing the specific facial features and vocalizations of each individual, chimpanzees are able to form social bonds and maintain a stable hierarchy within their group [57]. African elephants use olfactory and auditory cues for IR [33, 58]. They have a highly developed sense of smell and are able to recognize the unique scent of each member of their herd. Each elephant also has a distinct vocalization, which is used for communication within the herd. By recognizing the specific scent and vocalization of each individual, elephants are able to maintain social cohesion within their group and avoid aggressive interactions with unfamiliar elephants.

### 5.1 Conditions for a Signature System

We have derived a set of elementary conditions that are common across all signature systems, regardless of modality, including visual, acoustic, olfactory, and others. These conditions are critical for ensuring the accuracy and effectiveness of signature systems.

**Condition** (Universality). *Given a set of individuals in a given signature system, a signature of any individual must be necessarily and sufficiently unique, thus, universal*.

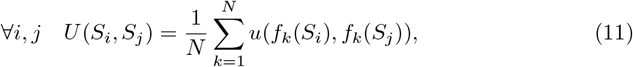

where *S*_*i*_ and *S*_*j*_ represent the signatures of two individuals, *N* ∈ ℕ_1_ represents the number of features being compared, *f*_*k*_ represents the *k*-th *feature function*, and *u* represents the comparison function that determines the similarity between the feature values for the two signatures. In this framing, the value of *U* (*S*_*i*_, *S*_*j*_) is a measure of the similarity between the signatures of two individuals. A lower value indicates a greater dissimilarity and hence a higher degree of universality between the signatures. The universality of an individual’s signature is determined by comparing their signature to those of other individuals and calculating the average similarity score.

Another formulation of this notion is: the ability of the signature system to maintain a low dissimilarity between signatures of the same individual across different *temporal points, conditions*, or *environments*. More formally, let 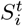 denote the signature of individual *i* at time *t*, and 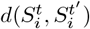 denote the *dissimilarity metric* between signatures of individual *i* at time *t* and time *t*^*′*^. The universality of the signature system for individual *i* can be defined as follows:

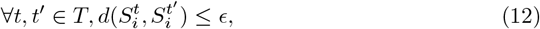

where *T* is the set of all possible time points, and *ϵ* ∈ ℝ^+^ is a small positive constant that represents the maximum allowable dissimilarity between signatures of the same individual across these temporal points. In other words, if the dissimilarity between signatures of the same individual exceeds the threshold *ϵ*, the signature system is deemed to have failed to maintain the universality of the individual’s signature. Therefore, a key challenge in biology or in artificial systems is designing signature systems is to minimize the dissimilarity between the signatures of the same individual while maximizing the dissimilarity between the signatures of different individuals. This ensures that the signature system can accurately recognize individuals while maintaining their universality over time, conditions, and environments.

**Condition** (Low Entropy). *For a given signature system, each signature must be entropy resistant and invariant to evolutionary pressures*.

To formulate the requirement for a signature to be entropy resistant and invariant to evolutionary pressures, we start by defining entropy as a measure of the amount of uncertainty or randomness in a system. Mathematically, this is represented by the equation:

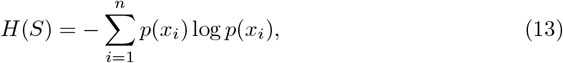

where *H*(*S*) is the entropy of the signature, *n* is the number of possible symbols in the signature, *p*(*x*_*i*_) is the probability of occurrence of symbol *x*_*i*_, and log is the logarithmic function. In the context of a signature system, we hypothesize that a signature must be entropy resistant, meaning that it should maintain a low entropy despite external or environmental pressures. This requires that the probability distribution of the signature’s symbols remain relatively constant, even when subjected to changes in the environment.

Furthermore, a signature must be invariant to evolutionary pressures, meaning that it must remain distinctive and easily distinguishable from others. Thus, we formulate the requirement for a signature to be entropy resistant and invariant to evolutionary pressures as:

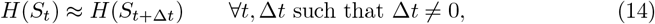

where *H*(*S*_*t*_) and *H*(*S*_*t*+Δ*t*_) represent the entropy of the signature at time *t* and time *t* + Δ*t* respectively, and Δ*t* represents a change in time. This formulation shows that the entropy of the signature must remain approximately constant over time. Thus, we think that a signature system’s requirement to be entropy resistant and invariance to evolutionary pressures emphasizes the importance of maintaining a low entropy signature that remains effective and distinct, even in the face of environmental and evolutionary changes. Low entropy signatures are important for IR in various species, such as primates, dolphins, and birds. These species use unique patterns of vocalizations, gestures, or physical markings as low entropy signatures for identification and communication within their social groups.

**Condition** (Unfalsifiability). *A signature is innately honest irrespective of the cost-benefit association to the signaler or the receiver*.

Let *S* be an individual’s signature, and let *E* be the set of all possible environments in which *S* can be verified. Let *F* (*S*) be the function that generates the features of *S*. Then, the unfalsifiability of *S* can be expressed as:

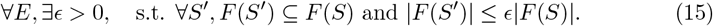

Eq (15) states that for any possible environment in which *S* may be verified, there exists a threshold value of similarity *ϵ* between *S* and any potential falsified signature *S*^*′*^, such that the features generated by *S*^*′*^ are a subset of the features generated by *S*, and the size of the feature set of *S*^*′*^ is less than or equal to a fraction of the size of the feature set of *S*. In other words, any attempt to falsify an individual’s signature must produce a signature that is so similar to the original that it cannot be distinguished from the original by the features used for verification.

**Condition** (Uniform Convergence). *Given a signature system, entities in it, under similar evolutionary pressures will converge to a stable state signature signaling equilibrium*.

Let us denote the signature system by 𝒮. Suppose that there are *n* entities (individuals) in 𝒮, denoted by *e*_1_, *e*_2_, …, *e*_*n*_. Let the signature of entity *e*_*i*_ be denoted by *s*_*i*_. The evolutionary pressures on can 𝒮 be represented by a function *f* (𝒮) that maps 𝒮 to a set of evolutionary pressures. We formalize this notion as follows: given signature system 𝒮 and a function *f* (𝒮) representing the evolutionary pressures on 𝒮, if all entities in *S* are subject to similar evolutionary pressures, then the signature system *S* will converge to a stable state signature signaling equilibrium, denoted by *E*_𝒮_. That is,

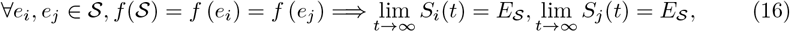

where *S*_*i*_(*t*) and *S*_*j*_(*t*) represent the signatures of entities *e*_*i*_ and *e*_*j*_ at time *t*, respectively. The stable state signature signaling equilibrium *E*_𝒮_ is a state in which all entities 𝒮 in have signatures that accurately distinguish them from other entities in 𝒮, while also being robust to environmental and other factors that may affect their signature characteristics. It ensures that small changes in the behavior or other factors do not lead to large changes in the signature. In addition, uniform convergence is related to the concept of stability, we address this relation in Section 6, which refers to the robustness of the signature with respect to noise or perturbations. A signature system that exhibits uniform convergence is generally more stable and less sensitive to small variations.

**Condition** (Cognitive Limit). *Cognitive Capacity is an upper bound on the complexity individuals in a signature system*.

Let 𝒮 be a signature system, and let *E* denote the set of entities in 𝒮. We define the cognitive capacity of an entity *i* ∈ *E* as *C*_*i*_, where *C*_*i*_ is a measure of the maximum complexity of signature that entity *i* can process and recognize in 𝒮. Let *S*_*i*_(*t*) denote the signature of entity *i* at time *t*, and let

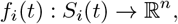

denote the mapping from the *signature space* of entity *i* to a feature space in ℝ^*n*^. We assume that the feature space is *shared* by all entities in 𝒮. We can then define the following equation for the cognitive capacity of entity *i*:

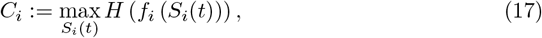

where *H* is the entropy function, and *f*_*i*_(*S*_*i*_(*t*)) is the feature representation of the signature *S*_*i*_(*t*). We hypothesize that the cognitive capacity of an entity *i* is an upper bound on the complexity of the signatures that entity *i* can process and recognize in 𝒮. That is, for any signature *S*_*i*_(*t*) that entity *i* encounters in 𝒮, the complexity of that signature cannot exceed *C*_*i*_.

## 6 Signature Systems are Attractor States

In this section, we attempt to connect IR with dynamical process modeling using attractor states. In the context of IR, attractor states are stable patterns of behavior or traits that become characteristic of a group or population, that can help distinguish individuals from each other based on *shared cues*. These patterns may arise from a variety of factors, such as social and environmental pressures, and can reinforce and shape the expression of individual identity over time.

Attractor states are often seen as emergent properties of *complex systems*, and can be modeled using mathematical approaches to understand their dynamics. *We hypothesize that individual identity cues in a signature system are attractor states*. This reinforcing state provides a basis for recognition and interaction. Individual identity cues can help shape the development of shared patterns of behavior that become attractor states.

To formally describe this connection between attractor states and individual identity cues, we start by defining a system of individuals, where each individual is represented by a set of characteristics or cues that distinguish them from others. We represent this system as *a set of vectors*, where each vector represents an individual and its associated cues. For example, if we have a system of honeybees, we might represent each honeybee as a vector with components for its size, coloration, scent, and other relevant characteristics:

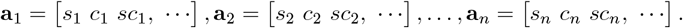

*s*_*i*_, *c*_*i*_, *sc*_*i*_, etc. represent the size, coloration, scent, and other relevant characteristics of individual *i*. We define attractor states as stable patterns of behavior or characteristics that emerge from interactions between individuals within the system. We represent these attractor states as *centroids or clusters* within the vector space defined by the individual cues. For example, we can define an attractor state for a group of individuals that tend to stay close together and move in unison:

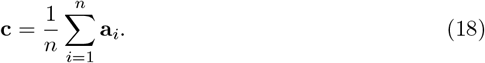

Here, **c** represents the centroid or center of mass for the set of individual vectors **a**_1_, **a**_2_, …, **a**_*n*_. This centroid can be seen as a representation of the *shared behavior or characteristics* that define the attractor state. We can describe the connection between individual identity cues and attractor states by noting that the cues can influence the formation and expression of these states. For example, if certain individuals within the system share similar cues, they may be more likely to interact with each other and develop shared patterns of behavior that become attractor states as illustrated in Fig. 5. Formally, we can represent this process as a clustering or grouping of individual vectors within the vector space defined by the cues. For example, we might use a clustering algorithm like K-Means to identify groups of individuals with similar coloration or scent cues:

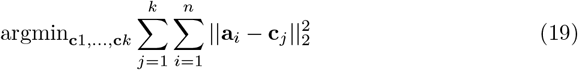

**Fig 5.**
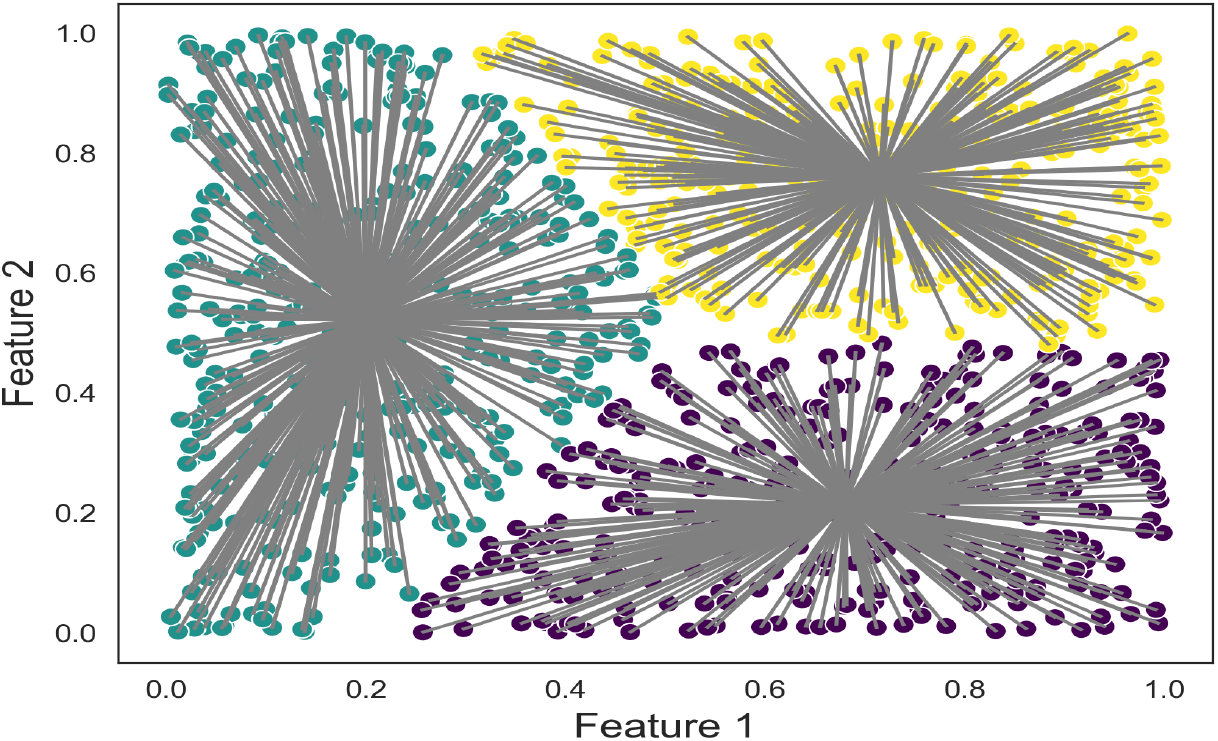
Scatter plot of individual vectors and cluster centroids obtained from K-means clustering of a set of uniform random individual vectors. The plot visualizes a two-dimensional data space consisting of 1000 individuals and 2 cues. The arrows pointing from the individual vectors towards the cluster centroids indicate the convergence of the individuals towards their respective attractor states.

Here, **c**_1_, …, **c**_*k*_ represent the centroids for *k* clusters of individual vectors, and the objective function minimizes the sum of squared distances between each individual vector and its nearest cluster centroid as shown in Eq (19).

This provides a way to formalize the relationship between individual identity cues and attractor states, and to analyze how these factors interact to shape the behavior and characteristics of individuals within a signature system.

Another way to look at this process is through dynamical steady-state theory where we show that signature systems can be reformulated as attractor states in a dynamical system. Let us define a signature system as a set of features or cues that are used to identify individuals. We represent each individual’s signature as a vector **s** ∈ ℝ^*n*^, where each component of the vector corresponds to the activity level of a particular feature or cue. We then define a function *f* : ℝ^*n*^→ [0, 1], which takes as input an individual’s signature **s** and produces as output a probability distribution over *possible identities*. This function is written as:

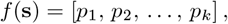

where *p*_*j*_ represents the probability that the individual corresponds to identity *j*. We use this function to define an attractor state in a dynamical system. Specifically, we define a dynamical system that evolves over time according to Eq (20):

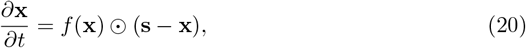

where **x** ∈ ℝ^*n*^ represents the state of the system at any given time, and ⊙ denotes element-wise multiplication (Hadamard product). This equation describes a system where each component of the state vector **x** is attracted towards the corresponding component of the individual’s signature **s** with a strength proportional to the probability that the individual corresponds to that identity. As time goes on, this system will tend to converge towards an attractor state where each component of the state vector is equal to the corresponding component of the individual’s signature. This attractor state represents the individual’s identity in the signature system as shown in Fig. 6. Thus, this form of representation shows that signature systems can be thought of as attractor states in a dynamical system.

**Fig 6.**
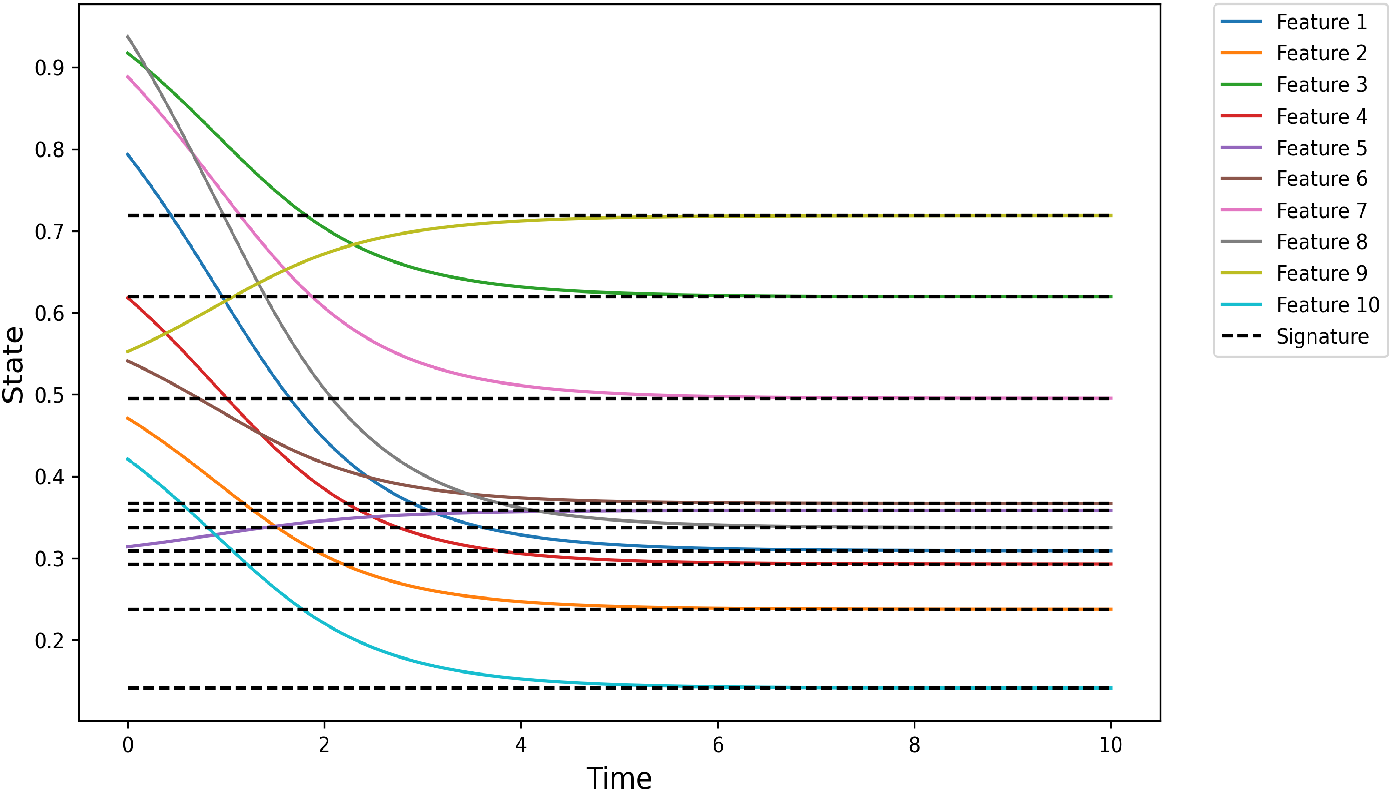
Simulation of a signature system as an attractor states in a dynamical system. The state of each feature or cue is plotted as a function of time, with the black dashed line representing the individual’s signature. The system converges towards the attractor state corresponding to the individual’s identity over time.

In this section, we have explored the link between individual identity cues and attractor states in a signature system. We hypothesize that individual identity cues in a signature system are attractor states that provide a basis for recognition and interaction. To describe this connection, we define a system of individuals represented by a set of cues that distinguish them from others. Attractor states emerge from interactions between individuals within the system and can be modeled as centroids or clusters within the vector space defined by the cues as shown in Fig. 5. These stable patterns of behavior or characteristics are influenced by individual identity cues and can shape the behavior and characteristics of individuals within a signature system. Using clustering algorithms like K-Means, we can analyze how individual identity cues interact with attractor states and shape shared patterns of behavior. We can also view signature systems as attractor states in a dynamical system, where each individual’s signature is represented as a vector, and the dynamics of the system lead to attractor states that provide a basis for individual recognition as shown in Figs. 6, 7, 8 and 9.

**Fig 7.**
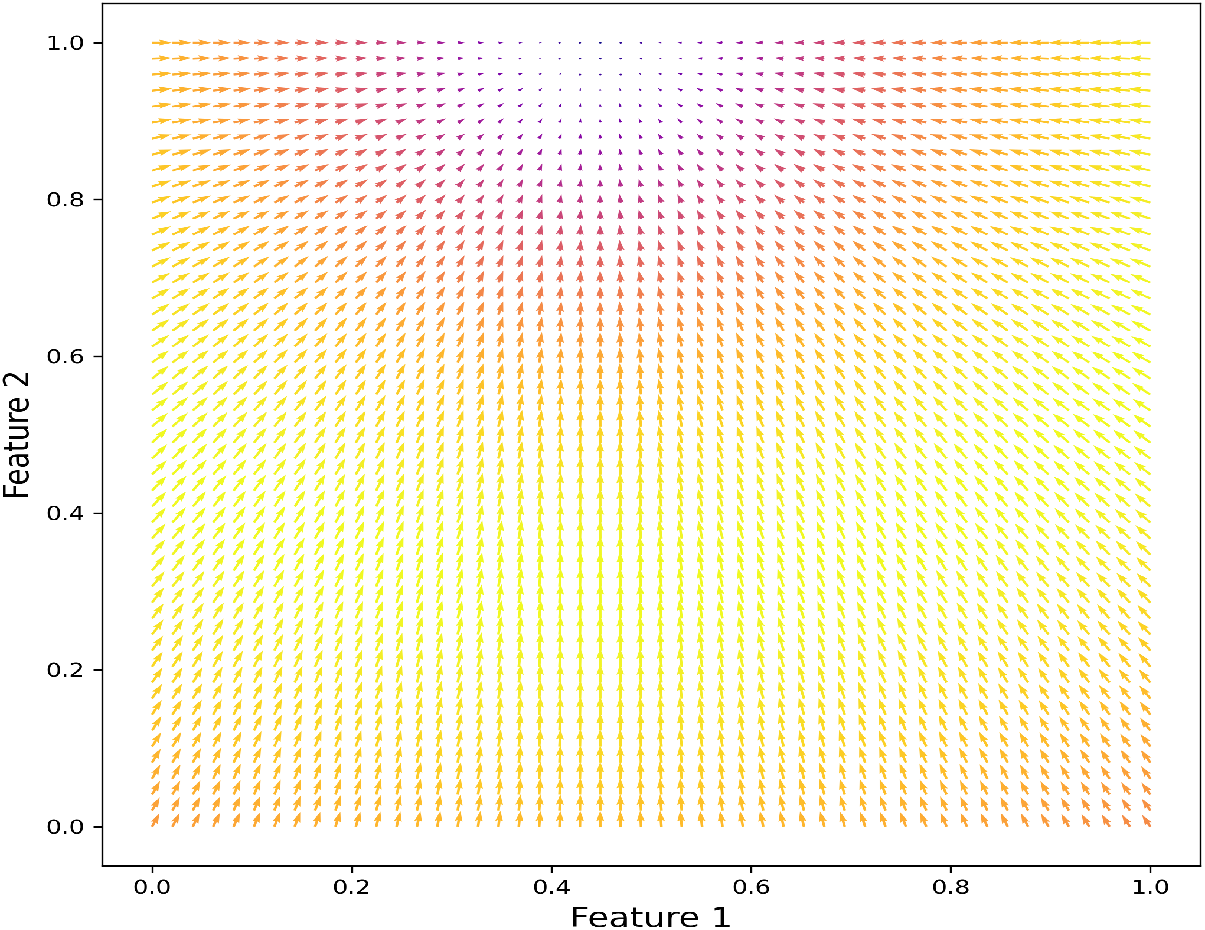
A vector field plot of the dynamical system, where the arrows indicate the direction and magnitude of the system’s evolution at each point in the 2D space. The color of the arrows is based on the magnitude of the vectors, with the warm colors (yellow and red) representing high magnitudes and the cool colors (blue and purple) representing low magnitudes.

**Fig 8.**
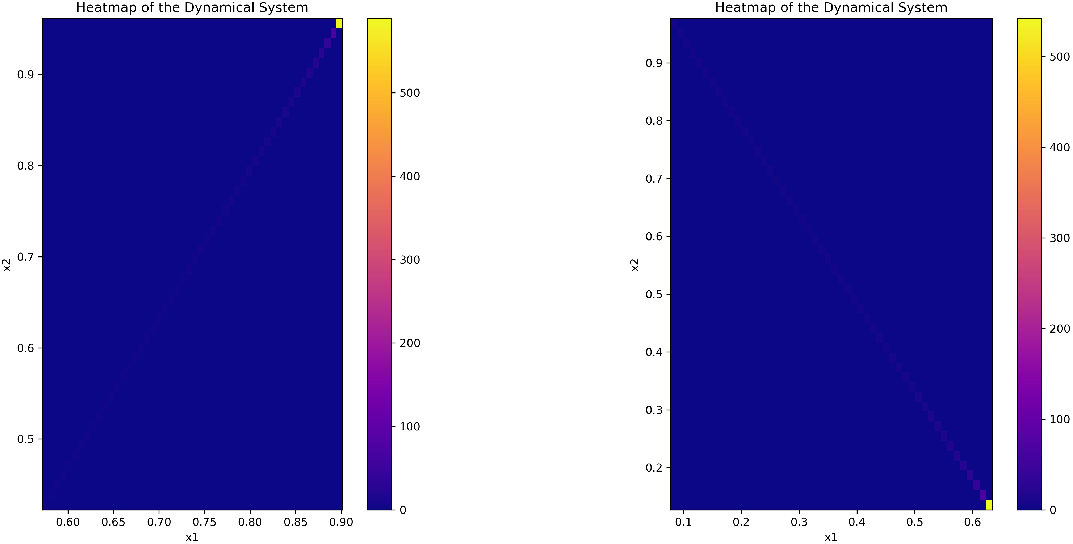
A heatmap plot of the probability of matching an individual’s signature in a two-dimensional feature space. The heatmap represents the probability values ranging from low (blue) to high (red) of an individual’s signature. The yellow dots in the plot show the initial and final positions of the dynamical system simulated over time.

**Fig 9.**
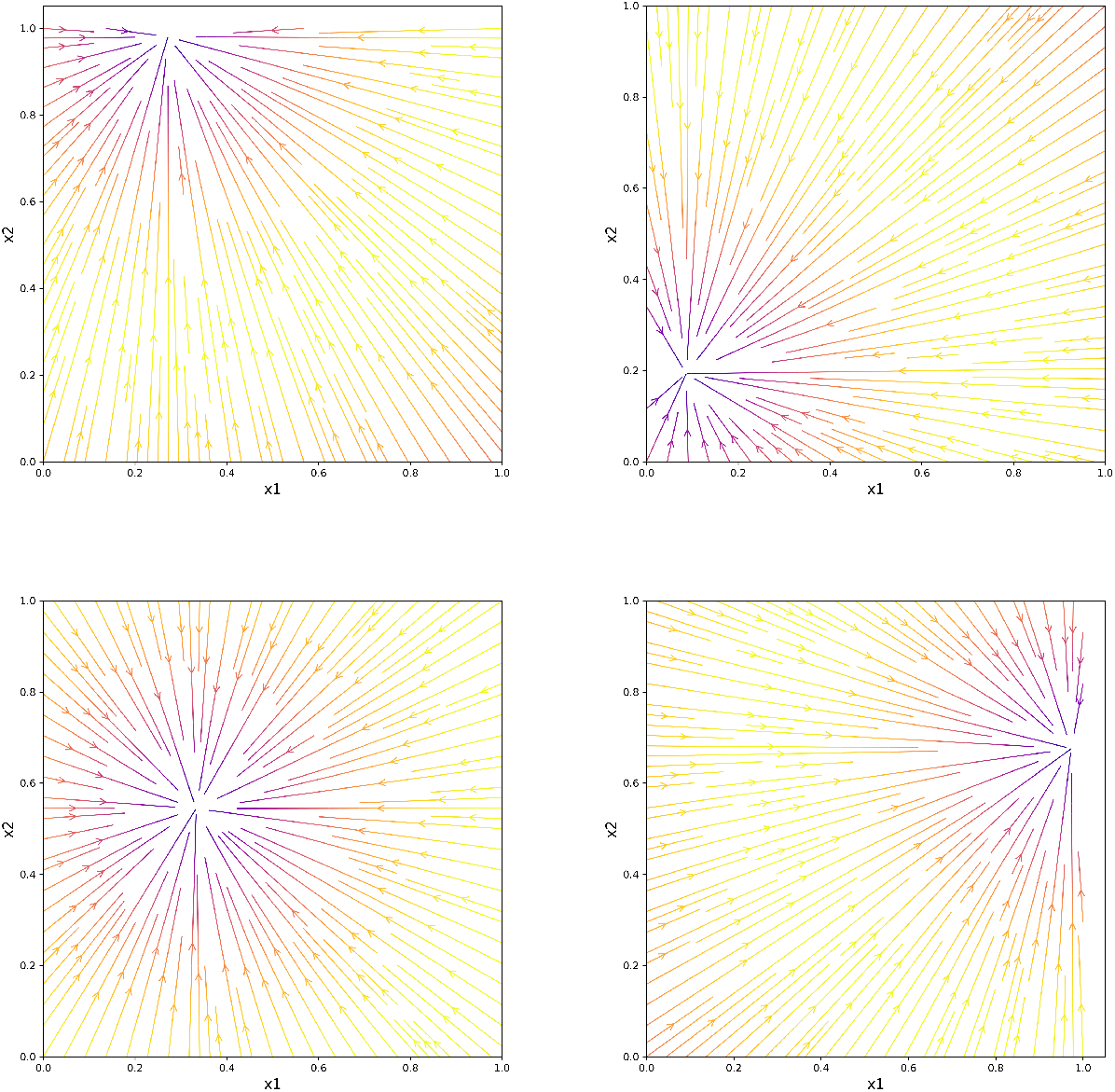
A phase portraits of a 2-dimensional dynamical system with an individual’s signature attraction field. The plot shows the trajectories of the system over time, starting from various initial conditions. The color of each trajectory represents the corresponding initial probability of matching the signature, with the warmer colors indicating higher probabilities.

## 7 Discussion

Recognition systems are vital for many social species, and understanding their underlying mechanisms can have fundamental and practical applications in varied fields such as conservation biology, animal behavior and communication, bio-metrics, etc. This paper presents a mathematical framework for recognition systems in animal communication and behavior. Our proposed framework has several potential applications. Firstly, we have made the first attempt, to the best of our knowledge, at formally defining individual recognition and disambiguate it from class-level recognition using principles from mathematics. Secondly, we proposed five conditions that must hold for true signature systems, ensuring the integrity of the underling signalling mechanism utilized for communication. In this paper, we strive to connect signature systems with dynamical systems modelling as biological processes are involve spatial and temporal dynamics. Our proposed framework for recognition systems has a significant potential for advancing our understanding of biological recognition and its broader implications for animal behavior and communication. IR involves complex cognition, which could help uncover more underlying principles of human language origins and evolution.

## 8 Conclusion

We present a theoretical framework for understanding biological recognition, utilizing a mathematical approach to identify the fundamental conditions of signature systems, such as the universality, low entropy, and unfalsifiability of an individual’s signature. The incorporation of category theory, information theory, dynamical systems modeling and optimization theory provide powerful tools for modeling recognition systems, applicable not only to biological systems but also to other fields, such as bio-metrics in computer science. We hope to motivate future research to focus on exploring the role of context in recognition systems and how recognition is affected by finer granularities that are manifested in real world scenarios. Technological advances may offer new opportunities for studying recognition systems at a cellular, molecular and neuronal levels. Overall, the mathematical conceptualization of IR provides a basis for advance our understanding of communication and the evolution of language.

## Acknowledgments

The project was financed by the funds of the research training group “Computational Cognition” (GRK2340) provided by the Deutsche Forschungsgemeinschaft (DFG), Germany

